# A minimal hybridization capture system for the parallel enrichment and cost-effective detection of ancient human pathogens

**DOI:** 10.1101/2025.06.02.657376

**Authors:** Arthur Kocher, Andaine Seguin-Orlando, Pierre Clavel, Guillaume Louvel, Richard Jonvel, Stéfan Tzortzis, Michel Signoli, Caroline Costedoat, Ludovic Orlando

## Abstract

The preservation of ancient DNA in archaeological remains enables identification of past disease agents. However, pathogen DNA is typically highly diluted by host and environmental DNA, limiting detection. Here, we present a proof-of-concept study using in-solution hybridization capture to improve detectability of a pre-defined set of 12 pathogens. We validate the method by detecting *Yersinia pestis*, the plague agent, in six individuals from 17^th^ and 18^th^ century French plague cemeteries with minimal sequencing. Expanding our probe set to target biomarkers of virtually any pathogen of interest offers a powerful tool for tracking the prevalence of infectious diseases in ancient populations.

## Background

Over the past two decades, significant advances in ancient DNA (aDNA) research have enabled the direct detection and genomic characterization of ancient pathogens whose genetic material may persist in skeletal and dental remains for thousands of years. These developments opened exciting avenues to reconstruct the epidemiological and evolutionary history of disease agents [1,2].

Early studies primarily relied on PCR-based assays targeting candidate pathogens responsible for historical epidemics (such as plague [3], and smallpox [4]), or those associated with paleopathological lesions in bone (e.g., Pott disease [5,6]). The development of high-throughput sequencing, alongside improved techniques for aDNA manipulation and computational identification, has since extended our capacity to characterize the metagenomic content of ancient biological samples [1,2,7]. As a result, genomic data have now been retrieved for approximately ∼50 obligatory or opportunistic pathogens dating back up to ∼10,000 years [8], primarily from human archaeological material, although ancient pathogens have also been identified in non-human animals [9–11], and plants [12,13]. These data sets offer unprecedented insights into the past genetic composition, phylogeographic trajectories, and functional evolution of these disease agents, including ancient lineages that predate their first appearance in historical records (e.g. the “Stone Age plague”, which predated by many thousand years the first plague pandemic known to the historical record [14,15]).

Despite these advances, ancient pathogen genomics continues to face several limitations. These include the patchy availability of archaeological collections, the low prevalence of infection among sampled individuals and differential DNA preservation in analyzed samples. Furthermore, pathogen DNA typically comprises only a tiny fraction of the total genetic material in infected remains, vastly outnumbered by host and environmental DNA, posing challenges for detection sensitivity [2,7,16]. These constraints help explain why ancient genomic data remains sparse for most pathogens, limiting our ability to characterize past microbial genetic diversity, and to make robust, quantitative inferences about past epidemiological dynamics.

While PCR amplification offers a rapid screening tool when pathogen-specific assays are available [17,18], it does not provide a comprehensive solution. Pathogens outside the targeted spectrum remain undetected, and the highly fragmented nature of aDNA often hinders amplification success. Moreover, issues such as non-specific amplification, contamination, limited sequence resolution, and the inability to genetically characterize the host, all constrain the utility and scope of PCR-based methods [19,20]. In contrast, shotgun sequencing has proven effective for recovering genome-wide sequence data from both pathogens and their hosts (e.g. [14,21,22]). However, this approach is resource-intensive, requiring high sequencing efforts and substantial post-processing, which may be prohibitively expensive or technically demanding for many laboratories.

Hybridization capture approaches may represent more cost-effective alternatives by concentrating sequencing efforts on organisms and genomic regions of interest [1,2]. These techniques have now been used extensively for the low-cost generation of ancient human genotyping data (e.g. [23]), as well as genome-wide data of ancient pathogens following initial detection through shallow shotgun sequencing [24–27]. However, the systematic application of hybridization capture to increase ancient pathogen detection sensitivity has so far been limited, despite bearing great potential.

Early attempts of ancient pathogen detection through hybridization capture systems allowing for the parallel enrichment of multiple pathogen species have shown encouraging results—for example, the Lawrence Livermore Microbial Detection Array or the Ancient Pathogen Screening Array [28,29]. However, these approaches have not been used since their initial development a decade ago, calling for the design and assessment of new protocols leveraging up-to-date biomolecular technologies.

Here, we present the design and evaluation of an in-solution hybridization capture system targeting taxonomically informative genomic markers to enable the parallel identification of multiple human pathogen species (Fig. 1). Our system employs commercially available and customizable in-solution RNA baits (*myBaits*®; Daicel Arbor Biosciences), which have been shown to outperform capture systems based on DNA microarrays or in-solution DNA baits [30]. While our proof-of-concept focuses on 12 selected pathogens, the system is readily scalable to include additional species with minimal added experimental cost. We assess its performance using skeletal remains from individuals buried in 17^th^ and 18^th^ century CE burials associated with documented outbreaks of bubonic plague, who likely died from *Yersinia pestis* infection [18,31].

**Fig. 1:**
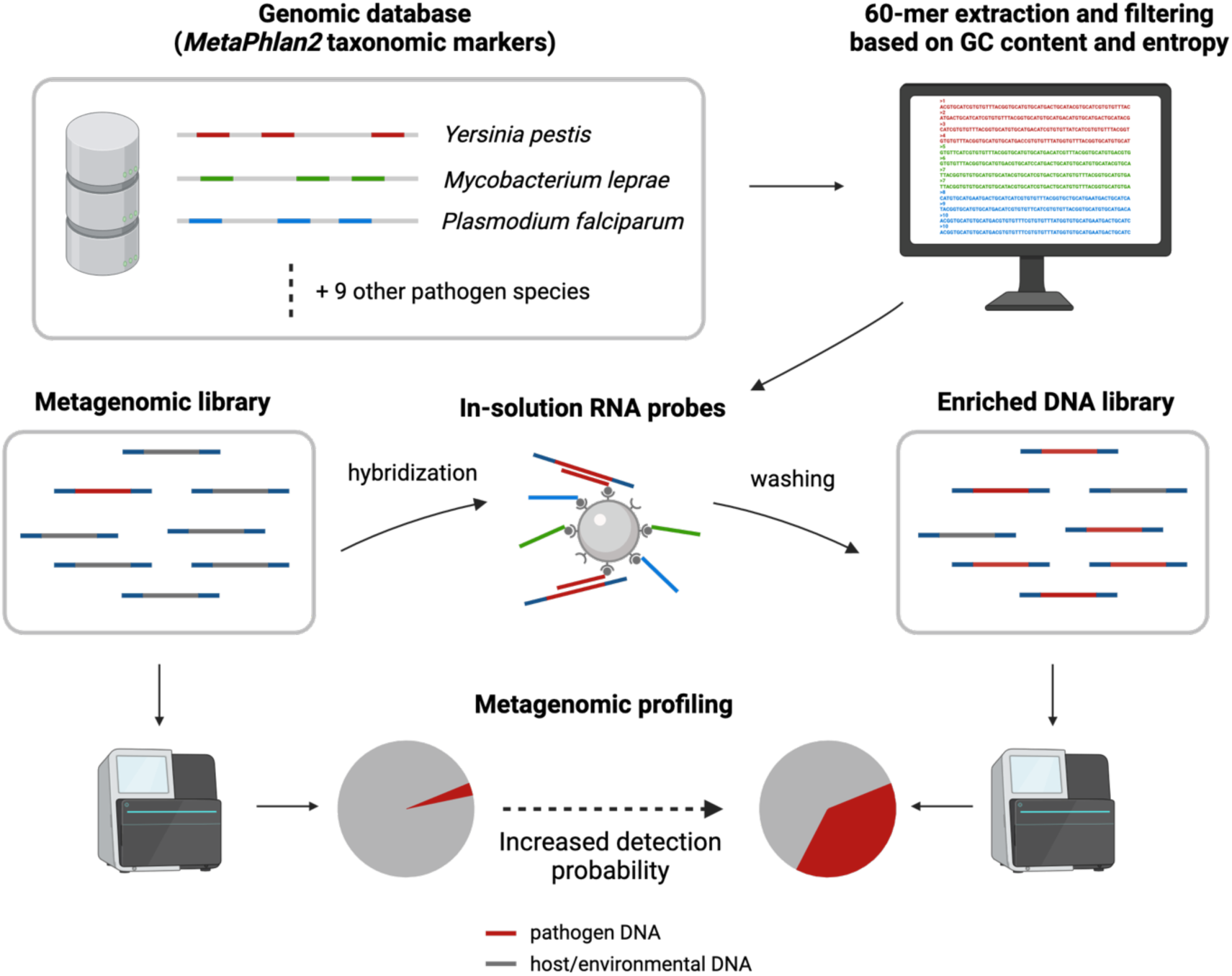
Schematic overview of the proposed hybridization capture system designed to enhance the detection probability of multiple pathogen species in parallel. Taxonomically informative markers from MetaPhlAn2 [32] are used to design pathogen-specific RNA probes for hybridization capture. Ancient DNA extracts are converted into amplified, immortalized DNA libraries. During hybridization, library fragments with sequence complementarity to the RNA probes form stable, transient complexes, while non-target DNA is removed through washing. This enrichment process effectively concentrates sequencing efforts on pathogen DNA that may be present in the original extract, enhancing detection efficiency and reducing background noise.

## Results

### Capture probe set design and laboratory processing of test samples

We retrieved taxonomically informative markers (N=1,036,027) from MetaPhlAn2 [32], representing a panel of 12 major human pathogens. These were selected to include key candidates responsible for major historical epidemics, including plague, leprosy, tuberculosis, syphilis, cholera, smallpox and malaria (Fig. 2a,b). The selected DNA sequence markers were then computationally fragmented into overlapping 60-mers with a 60 base pair tiling offset. Following, random down-sampling and filtering based on %GC content, entropy and the absence of undetermined bases, a subset of 1,048 to 3,529 probes were retained per pathogen, for a total of 21,570 probes. This number aimed at matching the minimum capacity of the myBaits Custom DNA-Seq kits, keeping experimental costs to a minimum.

**Fig. 2:**
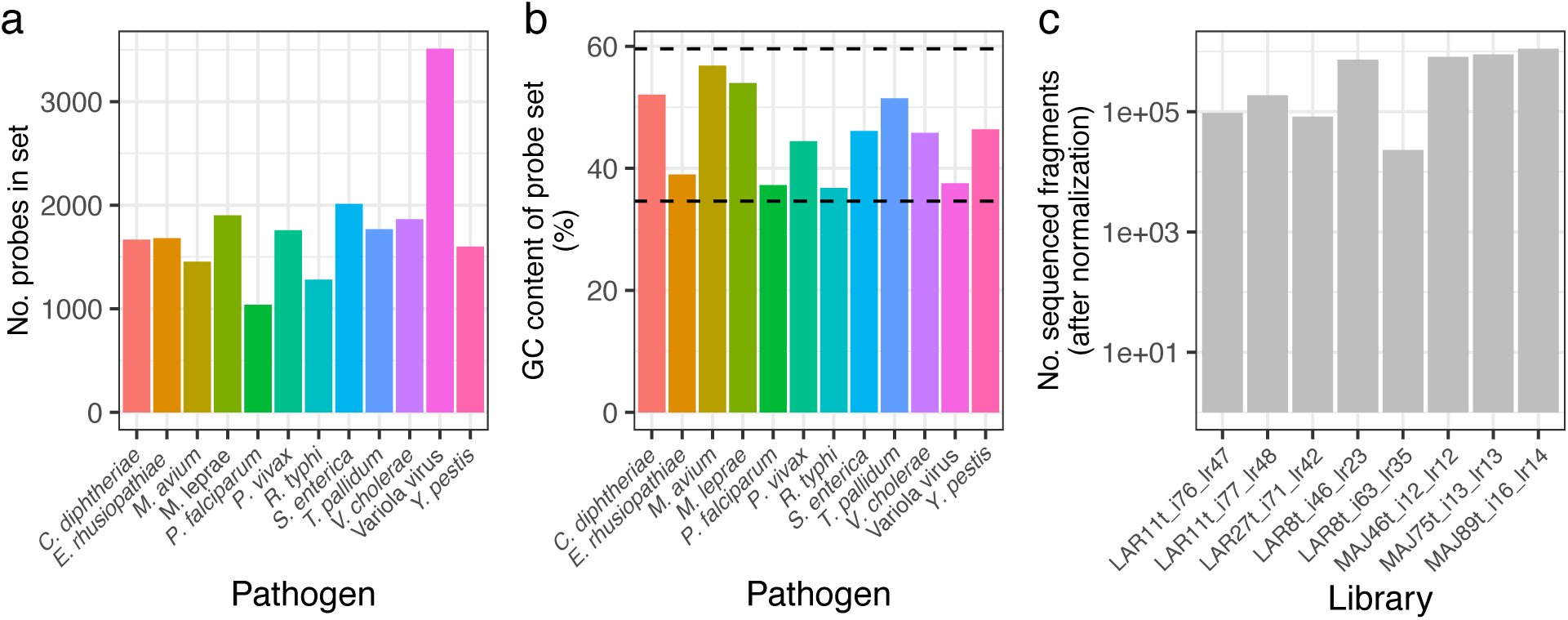
Probe set content and test sample sequencing depth. **a** Number of probes included in the capture system for each target pathogen, after filtering based on %GC content and entropy. **b** Average %GC content of the selected probes for each pathogen. Dotted lines indicate the %GC content range recommended to ensure stable hybridization with target sequences [33]. **c** Number of sequenced fragments for each test library, after pairwise normalization of shotgun and capture datasets.

To validate our hybridization capture assay, we prepared eight DNA libraries from the tooth remains of six individuals: three buried between 1629-1630 CE at the Lariey cemetery (Alps, France; samples LAR8t, LAR11t, and LAR27t), and three buried between 1720 and 1722 CE at the La Major site (Marseille, France; samples MAJ46t, MAJ75t, and MAJ89t) (Fig. 2c; Table 1). Both sites are associated with major plague epidemics in France: the former linked to outbreaks during the Thirty Years’ Wars, and the latter to the final plague occurrence of the second pandemics in the region [18,31]. Previous studies identified *Y. pestis*, the etiological agent of plague, in five of these individuals, through deep shotgun sequencing or following enrichment with a *Y. pestis*-specific capture system [18,31,34]. All DNA libraries were initially subjected to shallow shotgun sequencing for previous studies [18,31]. These same libraries were then enriched using our hybridization capture assay, generating an additional 0.02 to 2.15 million read pairs per individual. We then compared the post-capture data with the initial shotgun datasets, following normalization of sequencing efforts, to evaluate the performance of our system.

**Table 1:** Sequencing and mapping statistics of test samples.

| Library | No. reads after normalization | Prop. human in shotgun | No. YP reads* on target (shotgun/capture) | No. YP reads off target (shotgun/capture) | Clonality after capture (on/off-target) | Fold-enrichment (on/off-target) | Fold-enrichment after dedup. (on/off-target) |
| --- | --- | --- | --- | --- | --- | --- | --- |
| blanks (merged) | 309160 | 0 | 0 / 0 | 0 / 1 | 0 / 0 | Na / Na | Na / Na |
| LAR11t_i76_lr47 | 95153 | 0.043 | 16 / 33713 | 891 / 299 | 0.58 / 0.17 | 2107 / 0.34 | 891 / 0.28 |
| LAR11t_i77_lr48 | 187286 | 0.039 | 45 / 69437 | 1622 / 418 | 0.66 / 0.2 | 1543 / 0.26 | 525 / 0.21 |
| LAR27t_i71_lr42 | 82391 | 0.067 | 0 / 140 | 3 / 0 | 0.38 / 0 | >140 / 0 | >87 / 0 |
| LAR8t_i46_lr23 | 733334 | 0.169 | 28 / 52129 | 1223 / 520 | 0.8 / 0.47 | 1861 / 0.43 | 370 / 0.23 |
| LAR8t_i63_lr35 | 23015 | 0.168 | 0 / 1364 | 42 / 19 | 0.2 / 0.11 | >1374 / 0.45 | >1092 / 0.4 |
| MAJ46t_i12_lr12 | 818011 | 0.005 | 1 / 2246 | 20 / 14 | 0.97 / 0.57 | 2246 / 0.7 | 77 / 0.3 |
| MAJ75t_i13_lr13 | 894010 | 0.006 | 0 / 945 | 8 / 8 | 0.96 / 0.5 | >945 / 1 | 36 / 0.5 |
| MAJ89t_i16_lr14 | 1115723 | 0.004 | 0 / 10 | 1 / 0 | 0.5 / 0 | >10 / 0 | >5 / 0 |

\*High-quality alignments to the *Y. pestis* CO92 reference genome

### Y. pestis DNA enrichment efficiency and impact on screening results

Only one data set out of ten generated from experimental blanks (i.e., DNA libraries prepared from extracts performed without adding tooth sample powder) contained one single read mapping against the *Y. pestis* reference genome after hybridization capture (CO92 reference genome [35] ; Table 1). In contrast, all analyzed individuals associated with plague burials yielded at least one *Y. pestis* high-quality alignment following shotgun or hybridization capture, with significant variations consistent with previous results obtained from the same samples [18,31]. The number of high-quality alignments was sufficient to assess patterns of ancient DNA damage in only two of the libraries following shotgun sequencing, versus four following hybridization capture [7] (Fig. S1). Evidence of post-mortem deamination was limited, as reported previously for these individuals [31], and due to the treatment of DNA extracts by the USER enzymatic mix aimed at eliminating underlying signatures [36]. Base compositional profiles were, however, marked by a substantial over-representation of Cytosine (Guanine) residues at the reference position immediately flanking read starts (ends). This confirmed successful cleavage of Cytosine residues that were deaminated post-mortem [1], and authenticates the data generated.

Pairwise comparisons of shotgun and capture datasets revealed a significant increase in the number of *Y. pestis*-mapping after capture in all cases (Table 1; paired Wilcoxon Signed-Rank test, *p*-value = 0.004). This corresponds to an average enrichment of 54.7-fold (range=10.0-119.1 fold). Enrichment was especially marked when considering the genomic regions targeted by the capture probes relative to other genomic regions, which were depleted (Fig. 3ab), reaching an average of 1,939.5 (range=1,543-2,246) fold. The clonality of *Y. pestis* alignments was significantly increased following hybridization capture (Fig. 3c; paired Wilcoxon Signed-Rank test, *p*-value = 0.004), with an average of 63% (20%-97%) on target genomic regions, indicating that capture allowed recovering most of the targeted genomic material contained in the libraries even with shallow sequencing effort. Remarkably, the clonality of *Y.pestis* alignments falling outside of target regions also appeared increased (paired Wilcoxon Signed-Rank test, *p*-value = 0.016). It may reflect a combination of enrichment outside of targeted regions due to daisy chaining [31,33], and the loss of library complexity due to the additional amplification cycles performed to obtain sufficient material for capture.

**Fig. 3:**
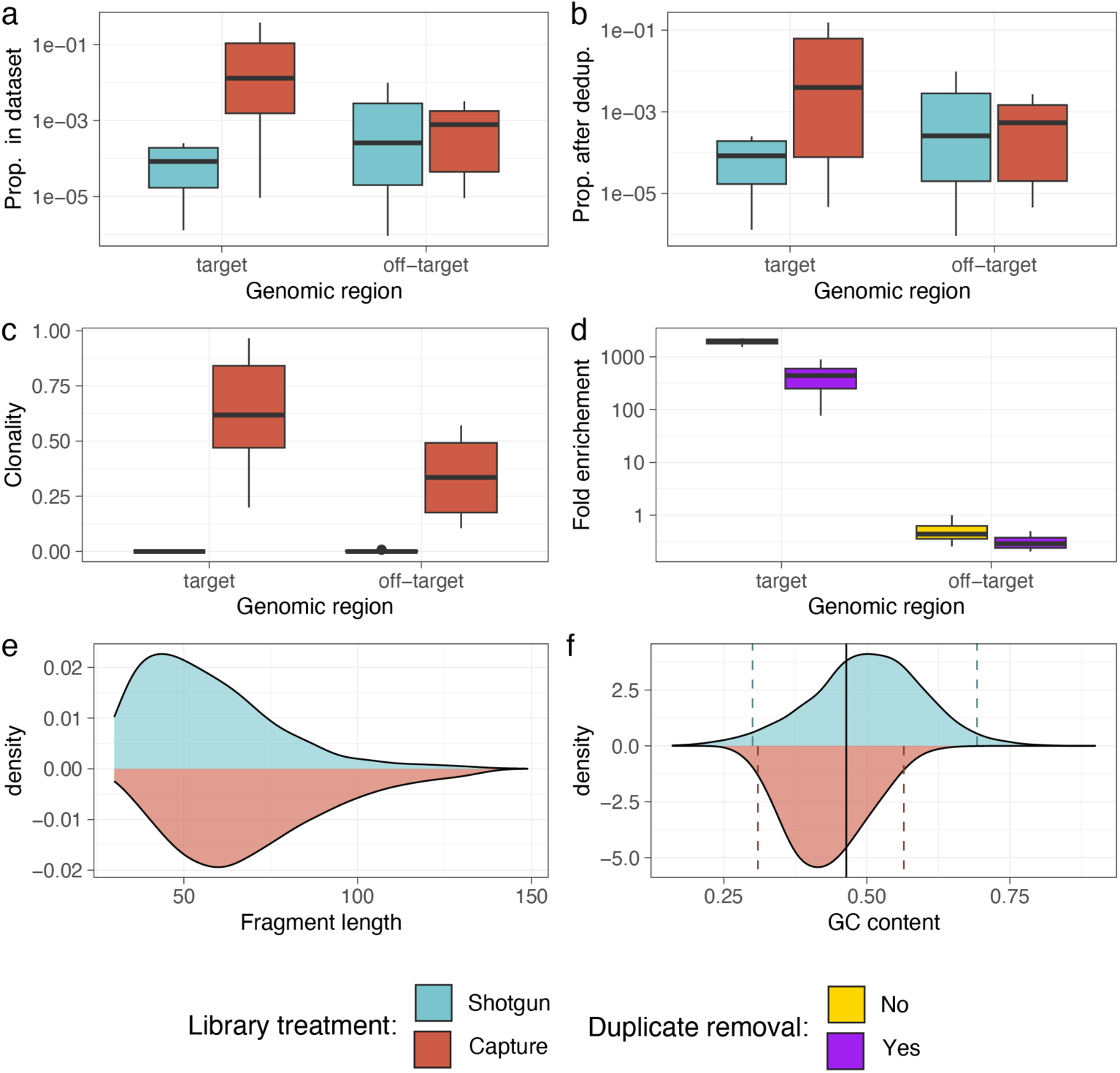
Assessment of capture efficiency. All shotgun and capture datasets were normalized pairwise for each library and mapped against the *Y. pestis* CO92 reference genome. Different metrics were derived from high quality *Y. pestis* alignments to assess the performance of the capture system and its impact on library content. **a** Boxplots of the fraction of *Y. pestis* fragments before and after capture, on target and off-target genomic regions. **b** Boxplots of the fraction of unique *Y. pestis* fragments. **c** Clonality of *Y. pestis* fragments before and after capture, on target and off-target genomic regions. **d** Boxplots of *Y. pestis* DNA fold-enrichment in target and off-target genomic regions. **e**. Density plot showing the distribution of *Y. pestis* fragment length before and after capture. **f** Density plot showing the distribution of %GC content in *Y. pestis* fragments before and after capture. The solid black line indicates the mean %GC content of the capture probes targeting *Y. pestis.* Colored dashed lines indicate 95% highest density intervals of the distributions.

Enrichment appeared significantly biased towards longer DNA fragments (Kolmogorov-Smirnov test, *p*-values < 1e-15), with a median length of high-quality alignments of 53 and 66 base pairs before and after hybridization capture, respectively (Fig. 3E), in line with previous work on hybridization capture [31,37]. This size shift likely results from the improved stability of probe-template dimers when templates can anneal to the whole probe RNA strand. Similarly, the base composition (%GC content) of high-quality alignments post-hybridization capture reflects that of the probe set, as expected for following successful enrichment. It was indeed significantly impacted post-hybridization capture (Kolmogorov-Smirnov test, *p*-values < 1e-15), and showed a shrunken distribution around the base composition of the probe set, as expected for following successful enrichment.

The analyses above measured the capture efficiency through raw comparisons of high-quality alignments against the *Y. pestis* genome, all of which are expected to hybridize preferentially to the capture probes whatever their actual taxonomic origin. However, the accurate detection of ancient microbial species in metagenomic datasets requires more specific assignment methods, including formal taxonomic assignment procedures [7].

To this end, we first performed metagenomic profiling of the datasets using the *metaBIT* pipeline [38]. Using a minimum relative abundance threshold of 1%, *Y. pestis* was detectable in three out of 8 of the shotgun datasets (Fig. 4a). In contrast, it became the dominant bacterial species in all libraries after capture, with the exception of MAJ89t_i16_lr14 for which no species could be detected.

**Fig. 4.**
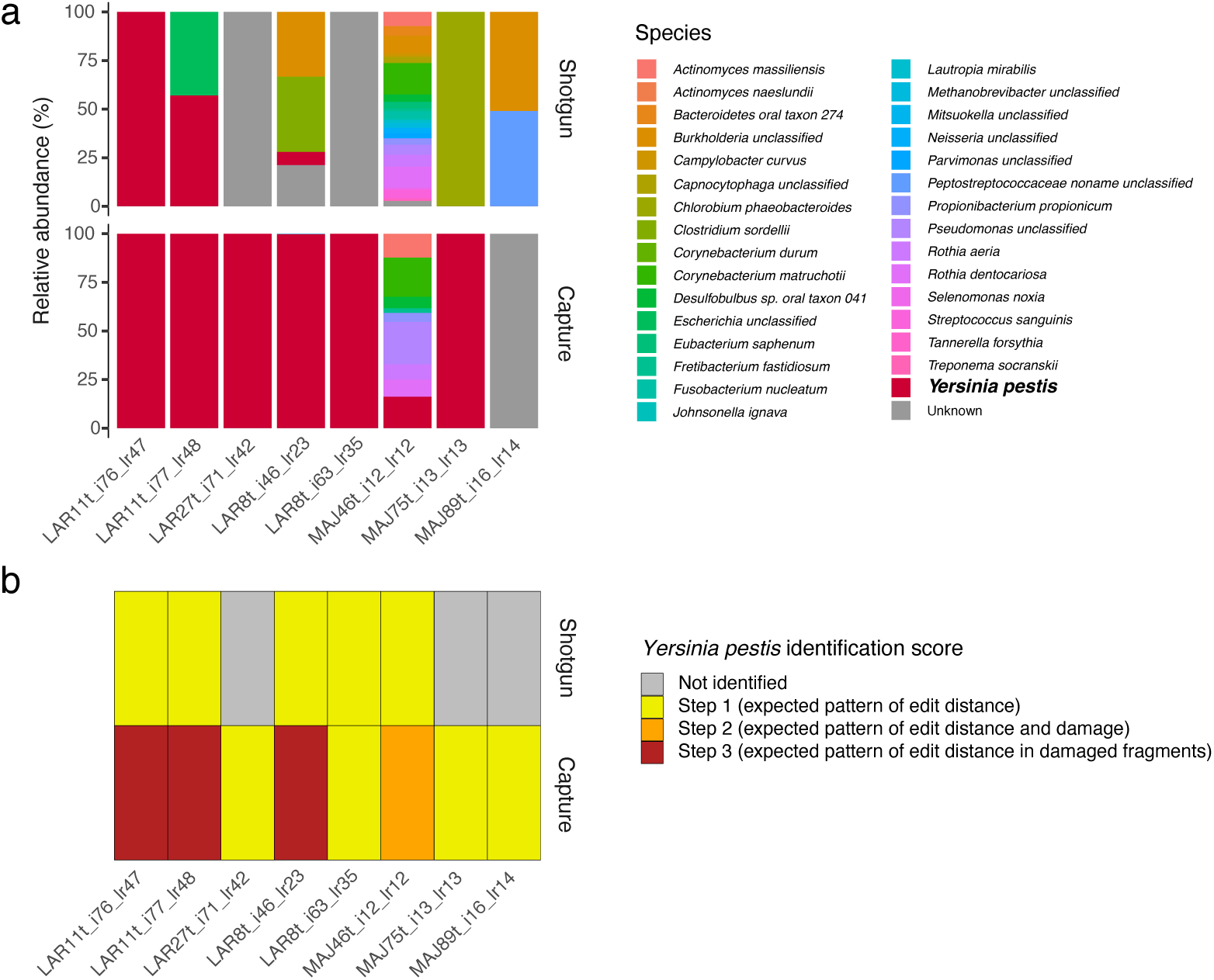
Metagenomic profiling and pathogen screening. **a** Results of metagenomic profiling with the *metaBIT* pipeline [38]. *Y. pestis* becomes the dominant bacterial species in most libraries following hybridization capture. **b** Results of *Y. pestis* detection using the *HOPS* ancient pathogen screening pipeline [39].

To further assess the authenticity of *Y. pestis* findings, we screened our datasets using the *HOPS* ancient pathogen pipeline [39] with an index containing one representative of all bacterial species included in the NCBI RefSeq database. The latter confirmed the presence of *Y. pestis* in 5/8 shotgun-sequenced libraries with the minimum identification step of 1 (indicating expected patterns of edit distance; Fig. 4). After capture, all eight libraries were positive for *Y. pestis*, and four of them exhibited identification scores > 1 indicating signatures of ancient DNA damage in assigned reads, despite minimal deamination patterns due to USER treatment (Fig. S1).

## Discussion

Identifying the etiological agents responsible for past epidemics remains challenging in the absence of genetic evidence confirming the presence of specific pathogens. Post-mortem molecular degradation significantly limits aDNA preservation and recovery, reducing the power of genetic identification. Additionally, many pathogens exhibit a tropism for soft tissues —which are rarely preserved—making detection impossible unless pathogens form lesions in the calcified skeletal and dental remains typically recovered during archaeological excavations, or penetrate the overall blood circulation.

Although highly specific, PCR-based detection assays often yield a high rate of false negatives when applied to the highly fragmented DNA typical of ancient samples [40]. Each assay also consumes a considerable portion of the aDNA extract while returning only limited sequence information, which represents a serious issue when extraction is destructive and sample material is scarce. By contrast, shotgun sequencing is perhaps the most comprehensive and conservative approach, offering a relatively unbiased snapshot of all DNA templates present in a sample. However, even this method is subject to biases introduced during library preparation and amplification, which can distort base composition and fragment size distributions [41]. More critically, shotgun sequencing is resource-intensive, as pathogen DNA often represents only a minute fraction of the total metagenomic content, requiring deep sequencing to detect target organisms reliably.

Hybridization capture provides a practical compromise between these two approaches. It combines the specificity of targeted detection with reduced sequencing demands, making it a cost-effective and efficient alternative. In this study, we have established that a simple hybridization capture assay can significantly improve the detection of pathogens, confirming the presence of *Y. pestis* in all six ancient individuals tested. In contrast, shotgun sequencing for an equivalent data could only identify three individuals positive for this infection (Fig. 4b). Excluding sequencing costs, the experimental hybridization procedure used in this study are associated with an estimated per-sample cost of approximately 100–170 euros, a figure that might vary across laboratories and can be further reduced by multiplexing a greater number of double-indexed DNA libraries (up to 23) during co-capture, as previously demonstrated [31,37].

Hybridization capture is also inherently scalable: the range of targeted pathogens can be expanded by increasing probe diversity, potentially encompassing the full spectrum of DNA viruses, bacteria, fungi, and parasites commonly associated with human infections. Moreover, the same capture assays can be adapted to recover genome-wide data from the host. Popular implementations typically target 1.3 million biallelic SNPs across the human genome, enabling comprehensive host genotyping [23]. This dual-target approach opens the door for integrated analyses of host-pathogen coevolution, offering insights into the genetic dynamics shaping immune responses over time. Provided sufficient sequence information is available, the system can also be applied to non-human organisms to explore pathogen loads across broader ecological and evolutionary contexts. Compared to shotgun sequencing, the increased sensitivity of hybridization capture is likely to yield more accurate estimates of pathogen prevalence in ancient populations. This, in turn, represents a critical step toward reconstructing past health landscapes and understanding how susceptibility to infectious diseases varied historically in response to environmental, sanitary, and social conditions.

The analyses presented in this study are based on a limited dataset, derived from only six individuals and minimal DNA sequence data. As these individuals originate from two relatively recent archaeological contexts in temperate environments, the performance of our hybridization capture system has not yet been evaluated under the wide range of preservation conditions and chronological depths represented in the archaeological record. Furthermore, our system could only be experimentally validated for *Y. pestis*, as this was the only pathogen detected in the tested individuals. However, since the RNA probes for all pathogens were designed using the same standardized approach, we can reasonably expect comparable performance for the additional targets included in the assay, pending further empirical validation.

## Conclusions

In this study, we developed a minimal probe set to enhance detection of 12 pathogens in ancient skeletal remains via hybridization capture. Our method identified the presence of *Y. pestis*, the plague agent, in all six individuals tested, despite limited sequencing efforts. This corresponds to a doubled detection rate compared to shotgun sequencing with similar effort. Our method relies on a simple probe design using extensive databases of microbial biomarkers that are publicly available. It can, thus, be readily expanded for the parallel detection of virtually any pathogen of interest, including DNA viruses as well as non-human pathogens, enabling targeted studies in animals and plants. This approach may be highly valuable for identifying the agents of past infectious diseases in the absence of diagnostic bone lesions and for assessing shifting prevalence rates across major historical transitions, enhancing our capacity to understand past sanitary conditions and their broader ecological and societal impacts.

## Methods

### Probe set design for the parallel enrichment of 12 human pathogens

We designed a custom *myBaits*® hybridization capture kit (Daicel Arbor Biosciences) to enrich taxonomically informative genomic regions in 12 human pathogens representing a broad spectrum of common infectious diseases: *Corynebacterium diphtheriae* (the agent of diphtheria), *Erysipelothrix rhusiopathiae* (erysipeloid), *Mycobacterium avium* (non-tuberculous mycobacteriosis), *Mycobacterium leprae* (leprosy), *Plasmodium falciparum* and *Plasmodium vivax* (malaria), *Rickettsia sp.* (typhus), *Salmonella enterica* (typhoid fever and other salmonelloses), *Treponema pallidum* (syphilis, yaws and bejel), variola virus (smallpox), *Vibrio cholerae* (Cholera), and *Yersinia pestis* (bubonic plague). For each of these except the variola virus, we designed hybridization probes targeting genomic regions included in the *MetaPhlAn2* reference database (v20_m200), which consists of clade-specific marker genes allowing unambiguous taxonomic assignments [32]. 60-mers were extracted from each marker gene, using a 60-bp tiling. For variola virus, 60-mers were extracted from the full reference genome (GenBank accession: NC_ 001611.1), using a 30-bp tiling. Unique probe sequences were then randomly sampled down to 2,140 for each pathogen, except for variola virus for which all 60-mers were retained. Only probes containing 34.6% to 59.6% GC, and showing entropy values ≥ 0.5427 were retained, to ensure stable hybridization with target sequences and sufficient sequence complexity [33]. Overall, we aimed for a total of ∼20,000 probes in the combined set after filtering, corresponding to the minimum capacity of the myBaits Custom DNA-Seq kits.

### Laboratory protocol and test samples

The efficiency of our capture system was evaluated on aDNA libraries prepared from the dental remains of six individuals excavated from graves associated with historic epidemics of bubonic plague in France. These included three individuals from the 1629-1630 CE plague cemetery of Lariey-Puy-Saint-Pierre in the French Alps (LAR), and three individuals from the site of La Major (MAJ) associated with the Great Plague of Marseille of 1720-1722 CE [18,31]. Given the context in which they were buried, all of these individuals were likely to have died from an infection with *Y. pestis.* This was previously confirmed for five of them based on Polymerase chain reaction (PCR) of a 133-bp fragment located on the *pla* gene (LAR8, LAR11 and LAR27; [18]), shotgun sequencing data (LAR8 and LAR11; [18]) or following enrichment using a *Y. pestis* whole-genome capture system (LAR27, MAJ46 and MAJ75; [31]).

Molecular work, from dental pulp sampling to sequencing library building and PCR setup preparation was carried out in the ancient DNA facilities of the CAGT (Université de Toulouse, France), following strict procedures to avoid and monitor contamination risks. Illumina sequencing libraries LAR11t_i76_lr47, LAR11t_i77_lr48, LAR8t_i63_lr35 and LAR27t_i71_lr42 were prepared for the study from Seguin-Orlando and colleagues [18], and libraries LAR8t_i46_lr23, MAJ46t_i12_lr12, MAJ75t_i13_lr13 and MAJ89t_i16_lr14 for the study from Clavel and colleagues [31]. Briefly, 40-70 mg of dental pulp was powdered, as described by Neumann and colleagues [42]. The DNA extraction was performed following a modified version of the Y2 protocol described by Gamba and colleagues [43], with a pre-digestion step of 1 hour at 37 °C, followed by a full digestion of the remaining pellet overnight at 42 °C. An aliquot of 200 μL of the overnight digestion lysate was purified on a MinElute column (QIAGEN©), and subjected to Uracil-Specific Excision Reagent (USER, NEB®) treatment. Sequencing libraries were constructed following a protocol described in [36] and modified as described in [44] to introduce a unique 7-nucleotide barcode within both adapters P5 and P7 (identified as *_lrXX* in the library name).

Libraries for shotgun sequencing were PCR enriched and indexed by performing 10-12 PCR cycles in 25 μL total reaction volume, using 1 unit of AccuPrimeTM Pfx DNA polymerase, 4 μL of unamplified DNA library and the InPE1.0 primer and one custom PCR primer (including a unique 6-nucleotide index (identified as *_iXX* in the library name), both at 200nM final concentration. Amplified products were purified using Agencourt Ampure XP beads (1.4:1 beads:DNA ratio) and eluted in 20 μL EB+0.05% tween.

Libraries for capture were PCR enriched and indexed as described in [31], by performing two consecutive rounds of PCR amplification. For the first amplification round, 8 parallel amplifications of 8 cycles were performed in a volume of 25 μL, using 2 μL of unamplified DNA library, 1 unit of AccuPrimeTM Pfx DNA polymerase and the InPE1.0 primer and one custom indexed PCR primer (200 nM final concentration). Products were pooled two by two, purified on four parallel MinElute columns (QIAGEN©), and eluted in 11 μL EB+0.05% tween each. Each of the four purified amplification products were split, and processed through two parallel PCR amplifications for 10 cycles, using for each PCR reaction 1 μL of the eluate in 25 μL total reaction volume. This second PCR round was carried out following the same conditions as for the first amplification round, except that IS5 and IS6 were used as PCR primers [45]. The resulting eight PCR products for each library were co-purified on a single MinElute column (QIAGEN©), and eluted in 12 μL EB+0.05% tween. Post-PCR library size profiles were checked on a Tapestation 4200 instrument (Agilent Technologies) and concentrations were measured on a QuBit HS dsDNA assay (Invitrogen). Before capture, the three MAJ libraries were pooled together, to obtain a final DNA quantity of 2.546 mg (0.412 mg of MAJ46t_i12_lr12, 1.052 mg of MAJ75t_i13_lr13 and 1.082 mg of MAJ89t_i16_lr14); while the five LAR libraries pooled together totalized 2.489 mg of DNA (0.548 for LAR11t_i76_lr47, 0.968 for LAR11t_i77_lr48, 0.031 mg for LAR8t_i63_lr35, 0.844 for LAR8t_i46_lr23 and 0.098 mg for LAR27t_i71_lr42). Pools were concentrated on a MinElute column down to a final volume of 7 μL. Capture was performed according to the myBaits® High Sensitivity Protocol Version 5.00, with two rounds of 18 to 22 hours hybridization with the probes at 55 °C. Captured libraries were PCR amplified for 14 cycles between both rounds, and for 8 cycles PCR after the second capture round (both using IS5 and IS6 primers and similar conditions as pre-capture PCR reactions).

Each final library, or library pool, was checked for size, molarity and concentration on a Tapestation 4200 instrument (Agilent Technologies) and on a QuBit HS dsDNA assay (Invitrogen) before being sequenced using the Paired-End mode on an Illumina MiniSeq instrument (2×81 cycles) at CAGT (Université de Toulouse, France), or on a NovaSeq 6000 (2×150 bp reads) at SciLifeLab (Stockholm, Sweden).

### Bioinformatic analyses and efficiency assessment

Sequencing read demultiplexing and collapsing, as well as adapter and poor-quality end trimming were performed using *AdapterRemoval2* v. 2.3.0 [46], tolerating at most one mismatch in each internal barcode (–barcode-mm-r[12] 1 –minadapteroverlap 3 –mm 5). Resulting merged and unmerged reads were concatenated for each dataset. The number of reads for shotgun and capture datasets were then equalized pairwise for each library by random downsampling using *SeqKit* v. 2.4.0 [47], to provide fair comparisons in subsequent analyses. To limit unspecific mapping to the *Y. pestis* genome, reads exhibiting low quality, low complexity or a length shorter than 30 bp were then removed using *fastp* v. 0.23.2 (using --length_required 30 --n_base_limit 3 --low_complexity_filter --trim_poly_g and default parameters otherwise).

All preprocessed datasets were then mapped against the human T2T-CHM13v2.0 reference genome [48] using *Bowtie 2* v. 2.5 [49] (with the --fast parameter setup), in order to compute the percentage of human fragments in each dataset, which were then filtered out for subsequent analyses. Remaining reads were mapped against the *Y. pestis* CO92 reference genome [35] using *Bowtie 2* v. 2.5 (with the --sensitive parameter setup), and alignments with a minimum mapping quality of 20 were extracted (*samtools view -c -F 4 -q 20*). PCR duplicates were removed using using MarkDuplicates from Picard Tools (version 2.18.7, http://broadinstitute.github.io/picard/). Damage patterns and base composition profiles of *Y. pestis* high-quality alignment (min. mapping quality of 20) were computed using mapDamage2 [50] and considering base with quality scores superior or equal to 30. Capture probe sequences were mapped against the *Y. pestis* CO92 reference genome following the same procedure to extract the genomic coordinates of the targeted regions using the *BEDTools* v. 2.31.1 (*bamtobed*, *sort* and *merge* commands).

To measure the enrichment efficiency of the hybridization capture system, we computed the ratio of the number of *Y. pestis-*mapping reads between pairwise capture and shotgun datasets. This enrichment factor was computed considering all genome positions globally, as well as for target and off-target regions separately (using the *samtools view -L* option with the *bed* file generated from the probe alignment as aforementioned). To assess the presence of an expected enrichment bias towards long and GC-rich DNA molecules, we further compared the fragment size and %GC content distribution of *Y. pestis*-mapping fragments in capture and shotgun datasets. Only merged reads with a length < 150 bp were considered for these comparisons, approximately corresponding to the maximum fragment length achievable with 2×81 cycles. Statistical significance of the observed differences was assessed using paired Wilcoxon Signed-Rank tests and Kolmogorov-Smirnov tests.

In order to assess the presence of *Y. pestis* DNA in the libraries more accurately, metagenomic profiling was performed using the *metaBIT* pipeline [38], restricting assignments to non-viral and non-eukaryotic taxa present in the MetaPhlAn2 database (v20_m200) [32]. Bacterial species supported by abundances lower than 1% were disregarded. Furthermore, we screened our datasets using the *HOPS* ancient pathogen pipeline [39] with an index containing one representative of all bacterial species included in the NCBI RefSeq database, using *Y. pestis-*mapping reads as input to reduce computational costs (with *Malt* parameters: *--minPercentIdentity 85, --topPercent 1, --gapOpen 1000, --maxAlignmentsPerQuery 100, --maxAlignmentsPerRef 10, --minSupport 1*).

## Declarations

### Ethics approval and consent to participate

All of the material and data used in this study derive from ancient DNA libraries generated in the context of previous studies and did not involve any extra destruction of archaeological material.

### Consent for publication

Not applicable

### Availability of data and materials

All sequencing data generated in the context of this study can be found on the European Nucleotide Archive (https://www.ebi.ac.uk/ena/browser/home) and are publicly available as of the date of publication under project accession number ENA: PRJEB89986.

### Authors’ contributions

LO designed the study. RJ, ST, MS, CC, and LO provided the study material. ASO and PC performed the experimental work. AK and GL performed the bioinformatic analysis. AK, ASO and LO wrote the manuscript.

## Acknowledgements

We thank Dr. Catherine Thèves, Dr. Clio Der Sarkissian and Pr. Norbert Telmon for their contributions to the original palaeogenomic studies involving some of the individuals analyzed here, Laure Calvière-Tonasso and Stéphanie Schiavinato for the CAGT ancient DNA facilities maintenance, as well as all the members of the AGES group at CAGT for fruitful discussion, and; Antoine Lacombe for hardware maintenance of the server cluster.

## Supplementary Information

**Supplementary Figure S1:**
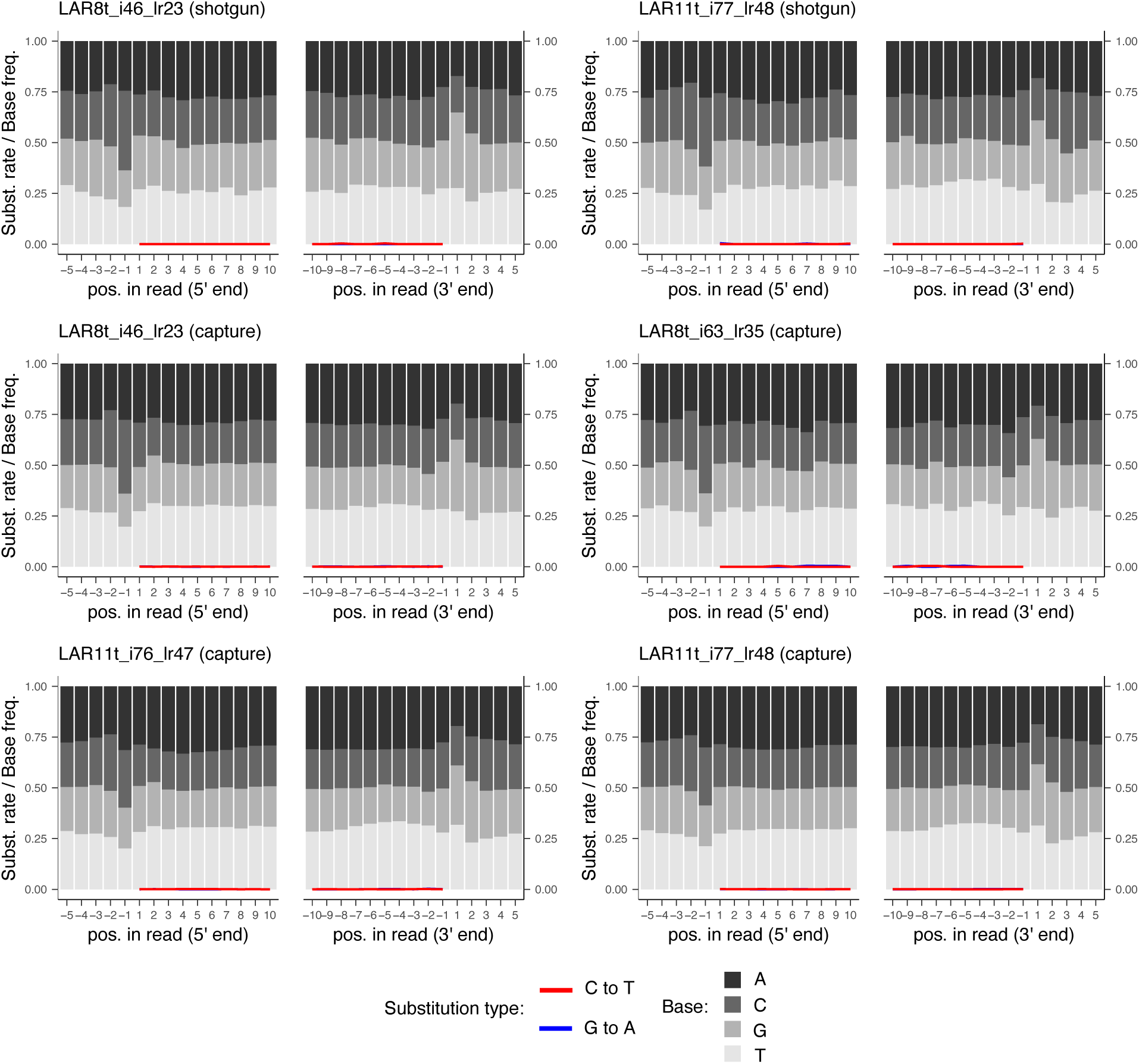
Damage patterns and base composition profiles of *Y. pestis* high-quality alignments following hybridization capture, for the four libraries yielding more than 1,000 unique alignments [7]. All aligned reads were processed using mapDamage2 [50] considering base with quality scores superior or equal to 30. Position-wise base composition profiles are provided for the 10 first (left, 1 to 10) and 10 last (right, -10 to -1) read positions within sequencing reads, and the 5 genomic positions preceding read starts (left, -5 to -1) or following read ends (right, 1 to 5). As the libraries were USER-treated, the DNA damage signature is an increase in %C at the alignment positions immediately preceding and following read termini [1].

